# Non-Vertical Cultural Transmission, Assortment, and the Evolution of Cooperation

**DOI:** 10.1101/2020.12.08.416560

**Authors:** Dor Cohen, Ohad Lewin-Epstein, Marcus W. Feldman, Yoav Ram

## Abstract

Cultural evolution of cooperation under vertical and non-vertical cultural transmission is studied, and conditions are found for fixation and coexistence of cooperation and defection. The evolution of cooperation is facilitated by its horizontal transmission and by an association between social interactions and horizontal transmission. The effect of oblique transmission depends on the horizontal transmission bias. Stable polymorphism of cooperation and defection can occur, and when it does, reduced association between social interactions and horizontal transmission evolves, which leads to a decreased frequency of cooperation and lower population mean fitness. The deterministic conditions are compared to outcomes of stochastic simulations of structured populations. Parallels are drawn with Hamilton’s rule incorporating assortment and effective relatedness.

## Introduction

Cooperative behavior can reduce an individual’s fitness and increase the fitness of its conspecifics or competitors [2]. Nevertheless, cooperative behavior appears to occur in many animals [5], including humans, primates [13], rats [24], birds [15, 27], and lizards [26]. Evolution of cooperative behavior has been an important focus of research in evolutionary theory since at least the 1930s [11, Appendix].

Since the work of Hamilton [12] and Axelrod & Hamilton [2], theories for the evolution of cooperative and altruistic behaviors have been intertwined often under the rubric of *kin selection*. Kin selection theory posits that natural selection is more likely to favor cooperation between more closely related individuals. The importance of *relatedness* to the evolution of cooperation and altruism was demonstrated by Hamilton [12], who showed that an allele that determines cooperative behavior will increase in frequency if the reproductive cost to the actor that cooperates, *c*, is less than the benefit to the recipient, *b*, times the relatedness, *r*, between the recipient and the actor. This condition is known as *Hamilton’s rule*:

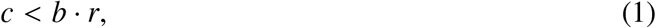

where the relatedness coefficient *r* measures the probability that an allele sampled from the cooperator is identical by descent to one at the same locus in the recipient.

Eshel & Cavalli-Sforza [6] studied a related model for the evolution of cooperative behavior. Their model included *assortative meeting*, or non-random encounters, where a fraction *m* of individuals in the population each interact specifically with an individual of the same phenotype, and a fraction 1 − *m* interacts with a randomly chosen individual. Such assortative meeting may be due, for example, to population structure or active partner choice. In their model, cooperative behavior can evolve if [6, eq. 3.2]

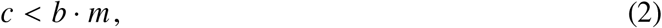

where *b* and *c* are the benefit and cost of cooperation^1^.

The role of assortment in the evolution of altruism was emphasized by Fletcher & Doebeli [9]. They found that in a *public-goods* game, altruism will evolve if cooperative individuals experience more cooperation, on average, than defecting individuals, and “thus, the evolution of altruism requires (positive) assortment between focal *cooperative* players and cooperative acts in their interaction environment.” With some change in parameters, this condition is summarized by [9, eq. 2.3]

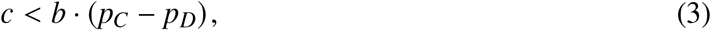

where *p_C_* is the probability that a cooperator receives help, and *p_D_* is the probability that a defector receives help.^2^ Bijma & Aanen [3] obtained a result related to inequality 3 for other types of games.

Cooperative behavior can be subject to *cultural transmission*, which allows an individual to acquire attitudes or behavioral traits from other individuals in its social group through imitation, learning, or other modes of communication [4, 25]. Feldman *et al.* [8] introduced the first model for the evolution of altruism by cultural transmission with kin selection and demonstrated that if the fidelity of cultural transmission of altruism is *φ*, then the condition for evolution of altruism in the case of sib-to-sib altruism is [8, Eq. 16]

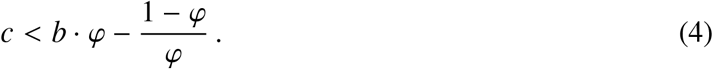

In inequality 4, *φ* takes the role of relatedness (*r* in inequality 1) or assortment (*m* in inequality 2), but the effective benefit *b*·*φ* is reduced by (1 − *φ*)/*φ*. This shows that under a cultural transmission, the condition for the evolutionary success of altruism entails a modification of Hamilton’s rule (inequality 1).

Cultural transmission may be modeled as vertical, horizontal, or oblique: vertical transmission occurs between parents and offspring, horizontal transmission occurs between individuals from the same generation, and oblique transmission occurs to offspring from the generation to which their parents belong (i.e. from non-parental adults). Evolution under either of these transmission models can be be more rapid than under pure vertical transmission [4, 21, 23]. Both Woodcock [28] and Lewin-Epstein *et al.* [16] demonstrated that non-vertical transmission can help explain the evolution of cooperative behavior, the former using simulations with cultural transmission, the latter using a model where cooperation is mediated by host-associated microbes. Indeed, models in which microbes affect their host’s behavior [10, 16, 17] are mathematically similar to models of cultural transmission, and they also emphasize the role of non-vertical transmission [4].

Here, we study models for the cultural evolution of cooperation that include both vertical and non-vertical transmission. In our models behavioral changes are mediated by cultural transmission that can occur specifically during social interactions. For instance, there may be an association between the choice of partner for social interaction and the choice of partner for cultural transmission, or when an individual interacts with an individual of a different phenotype, exposure to the latter may lead the former to convert its phenotype. Our results demonstrate that cultural transmission, when associated with social interactions, can enhance the evolution of cooperation even when genetic transmission cannot, partly because it can facilitate the generation of assortment [9], and partly because it can diminish the effect of natural selection [23]. This further emphasizes that treatment of cooperation as a cultural trait, rather than a genetic one, can lead to a broader understanding of its evolutionary dynamics.

## Models

Consider a large population whose members can be one of two phenotypes: *ϕ* = *A* for cooperators or *ϕ* = *B* for defectors. An offspring inherits its phenotype from its parent via vertical transmission with probability *v* or from a random individual in the parental population via oblique transmission with probability (1 − *v*) (Figure 1a). Following Ram *et al.* [23], given that the parent’s phenotype is *ϕ* and assuming uni-parental inheritance [29], the conditional probability that the phenotype *ϕ*′ of the offspring is *A* is

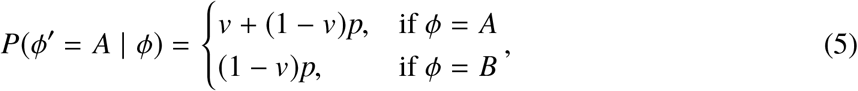

where *p* = *P*(*ϕ* = *A*) is the frequency of *A* among all adults in the parental generation.

**Figure 1:**
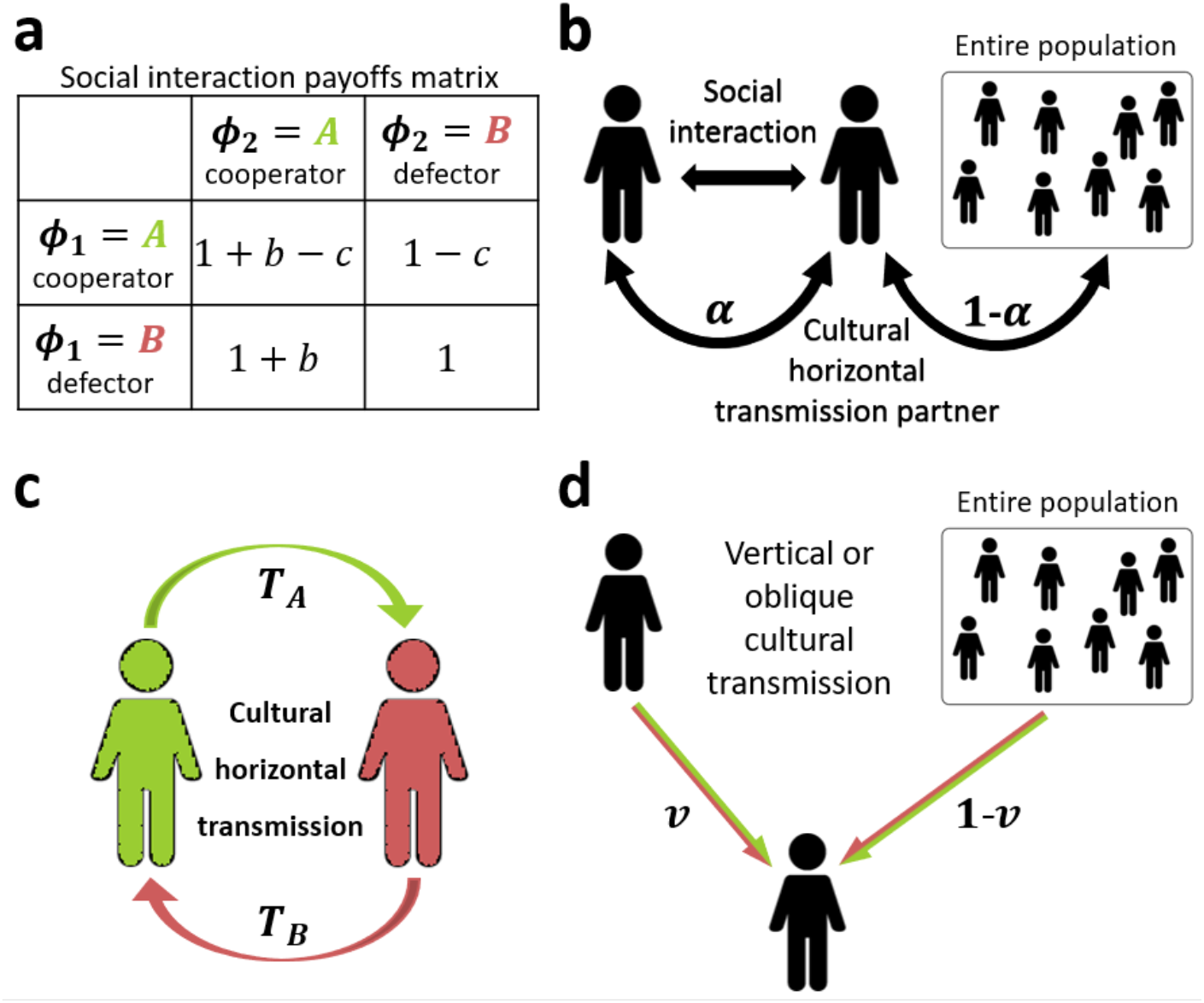
Model illustration. **(a)** Prisoners’ dilemma payoff matrix, showing The fitness of phenotype *ϕ*_1_ when interacting with phenotype *ϕ*_2_. **(b)** Individuals socially interact in pairs in a prisoners’ dilemma game. Horizontal cultural transmission occurs from a random individual in the population, with probability 1 − *α*, or from the social partner, with probability *α*, where *α* is the interaction-transmission association parameter. **(c)** The probabilities of successful horizontal cultural transmission of phenotypes *A* and *B* are *T_A_* and *T_B_*, respectively. **(d)** Offspring inherit their parent’s phenotype with probability *v*, or the phenotype of a random non-parental adult with probability 1 − *v*.

Not all adults become parents, and we denote the frequency of phenotype *A* among parents by 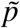. Therefore, the frequency 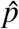 of phenotype *A* among juveniles (after selection and vertical and oblique transmission) is

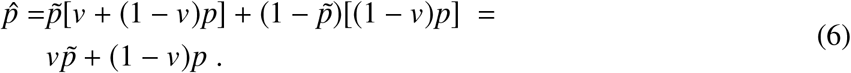

Individuals are assumed to interact according to a *prisoner’s dilemma*. Specifically, individuals interact in pairs; a cooperator suffers a fitness cost 0 < *c* < 1, and its partner gains a fitness benefit *b*, where we assume *c* < *b*. Figure 1a shows the payoff matrix, i.e. the fitness of an individual with phenotype *ϕ*_1_ when interacting with a partner of phenotype *ϕ*_2_.

Social interactions occur randomly: two juvenile individuals with phenotype *A* interact with probability 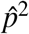, two juveniles with phenotype *B* interact with probability 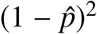, and two juveniles with different phenotypes interact with probability 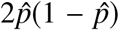. Horizontal cultural transmission occurs between pairs of individuals from the same generation. It occurs between socially interacting partners with probability *α*, or between a random pair with probability 1 − *α* (see Figure 1b). However, horizontal transmission is not always successful, as one partner may reject the other’s phenotype. The probability of successful horizontal transmission of phenotypes *A* and *B* are *T_A_* and *T_B_*, respectively (Table 1, Figure 1c). Then, the frequency *p*′ of phenotype *A* among adults in the next generation, after horizontal transmission, is

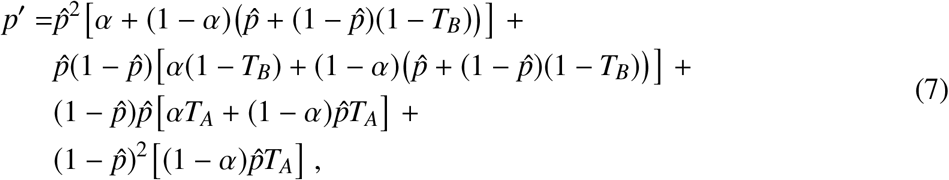

which simplifies to

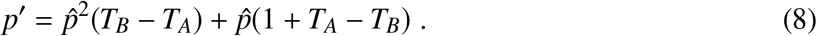

**Table 1:** Interaction frequency, fitness, and transmission probabilities.

| Phenotype $\phi_1$ | Phenotype $\phi_2$ | Frequency | Fitness of $\phi_1$ | $P(\phi_1 = A)$ via horizontal transmission: | |
| --- | --- | --- | --- | --- | --- |
| | | | | from partner, $\alpha$ | from population, $(1 - \alpha)$ |
| $A$ | $A$ | $\hat{p}^2$ | $1 + b - c$ | 1 | $\hat{p} + (1 - \hat{p})(1 - T_B)$ |
| $A$ | $B$ | $\hat{p}(1 - \hat{p})$ | $1 - c$ | $1 - T_B$ | $\hat{p} + (1 - \hat{p})(1 - T_B)$ |
| $B$ | $A$ | $\hat{p}(1 - \hat{p})$ | $1 + b$ | $T_A$ | $\hat{p}T_A$ |
| $B$ | $B$ | $(1 - \hat{p})^2$ | 1 | 0 | $\hat{p}T_A$ |

The frequency of *A* among parents (i.e. after selection) follows a similar dynamic, but also includes the effect of natural selection, and is therefore

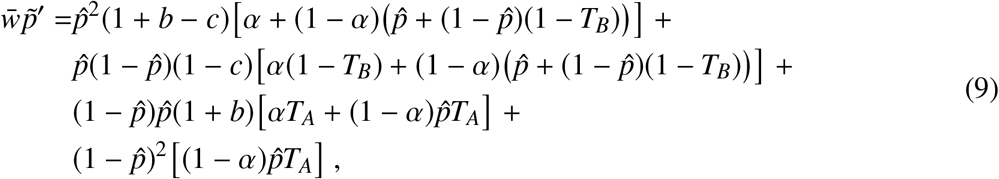

where fitness values are taken from Figure 1a and Table 1, and the population mean fitness is 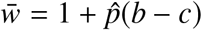. Eq. 9 can be simplified to

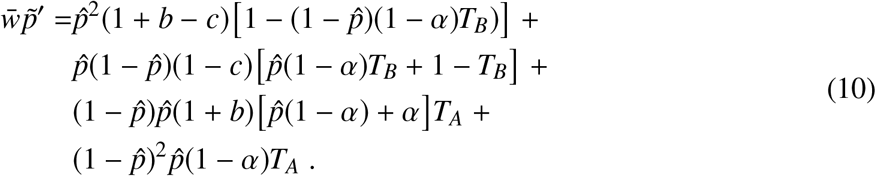

Starting from Eq. 6 with 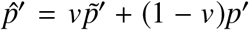, we substitute *p*′ from Eq. 8 and 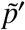 from Eq. 10 and obtain

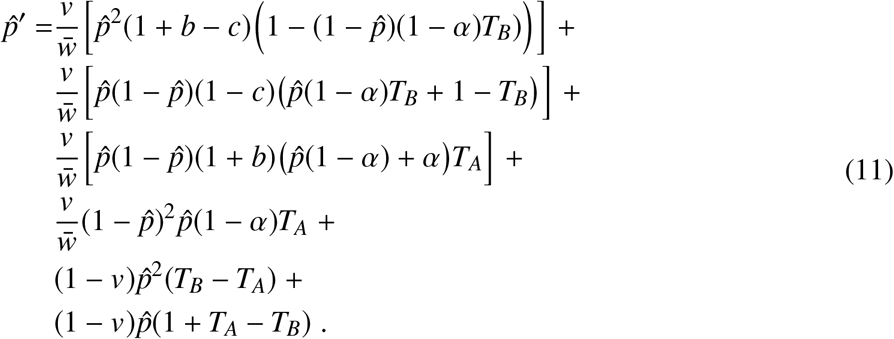

Table 2 lists the model variables and parameters.

**Table 2:** Model variables and parameters.

| Symbol | Description | Values |
| --- | --- | --- |
| $A$ | Cooperator phenotype | |
| $B$ | Defector phenotype | |
| $p$ | Frequency of phenotype $A$ among adults | $[0, 1]$ |
| $\tilde{p}$ | Frequency of phenotype $A$ among parents | $[0, 1]$ |
| $\hat{p}$ | Frequency of phenotype $A$ among juveniles | $[0, 1]$ |
| $v$ | Vertical transmission rate | $[0, 1]$ |
| $c$ | Cost of cooperation | $(0, 1)$ |
| $b$ | Benefit of cooperation | $c < b$ |
| $\alpha$ | Probability of interaction-transmission association | $[0, 1]$ |
| $T_A, T_B$ | Horizontal transmission rates of phenotype $A$ and $B$ | $(0, 1)$ |

## Results

We determine the equilibria of the model in Eq. 11, namely, solutions of 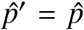, and analyze their local stability. We then analyze the evolution of a modifier of interaction-transmission association, *α*. Finally, we compare derived conditions to outcomes of stochastic simulations with a structured population.

### Evolution of cooperation

The fixed points (equilibria) of the recursion (Eq. 11) are 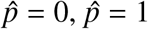, and (see Eq. B5)

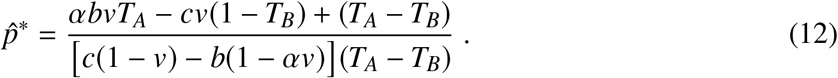

Define the following cost thresholds, *γ*_1_ and *γ*_2_, and the vertical transmission threshold, 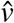,

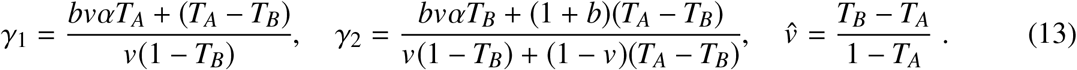

Then we have the following result.

#### Result 1 Equilibria and stability

*With vertical, horizontal, and oblique transmission, the cultural evolution of a cooperation follows one of the following scenarios in terms of the cost thresholds γ*_1_ *and γ*_2_ *and the vertical transmission threshold* 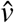 *(Eq. 13):*

1. Fixation of cooperation*: if* (i) *T_A_* ≥ *T_B_ and c* < *γ*_1_*; or if* (ii) *T_A_* < *T_B_ and* 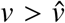 *and c* < *γ*_2_.
2. Fixation of defection*: if* (iii) *T_A_* ≥ *T_B_ and γ*_2_ < *c; or if* (iv) *T_A_* < *T_B_ and γ*_1_ < *c.*
3. Stable polymorphism*: if* (v) *T_A_* < *T_B_ and* 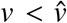 *and c* < *γ*_1_*; or if* (vi) *T_A_* < *T_B_ and* 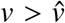 *and γ*_2_ < *c* < *γ*_1_.
4. Unstable polymorphism*: if* (vii) *T_A_* > *T_B_ and γ*_1_ < *c* < *γ*_2_.

These conditions are illustrated in Figures 2a, 2b, 3a, and 3b, and the full analysis is in Appendix B.

**Figure 2:**
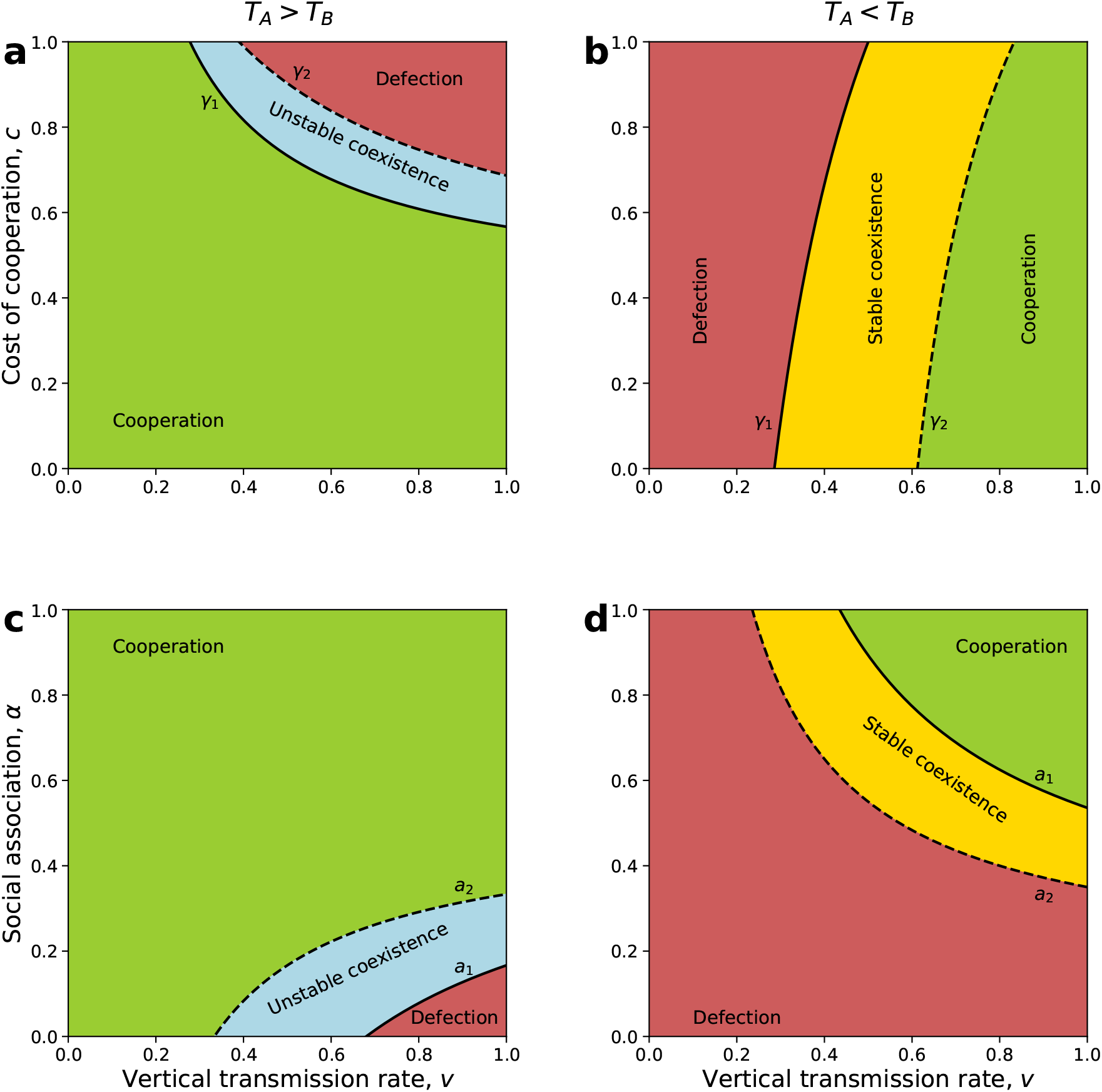
Evolution of cooperation under vertical, oblique, and horizontal cultural transmission. The figure shows parameter ranges for global fixation of cooperation (green), global fixation of defection (red), fixation of either cooperation or defection depending on the initial conditions, i.e. unstable polymorphism (blue), and stable polymorphism of cooperation and defection (yellow). In all cases the vertical transmission rate *v* is on the x-axis. (**a-b**) Cost of cooperation *c* is on the y-axis and the cost thresholds *γ*_1_ and *γ*_2_ (Eqs. 13) are represented by the solid and dashed lines, respectively. (**c-d**) Interaction-transmission association *α* is on the y-axis and the interaction-transmission association thresholds *a*_1_ and *a*_2_ (Eqs. 18) are represented by the solid and dashed lines, respectively. Horizontal transmission is biased in favor of cooperation, *T_A_* > *T_B_*, in (**a**) and (**c**), or defection, *T_A_* < *T_B_*, in (**b**) and (**d**). Here, *T_A_* = 0.5, and (**a**) *b* = 1.2,, *T_B_* = 0.4, *α* = 0.4; (**b**) *b* = 2, *T_B_* = 0.7, *α* = 0.7; (**c**) *b* = 1.2, *T_B_* = 0.4, *c* = 0.5; (**d**) *b* = 2, *T_B_* = 0.7, *c* = 0.5.

**Figure 3:**
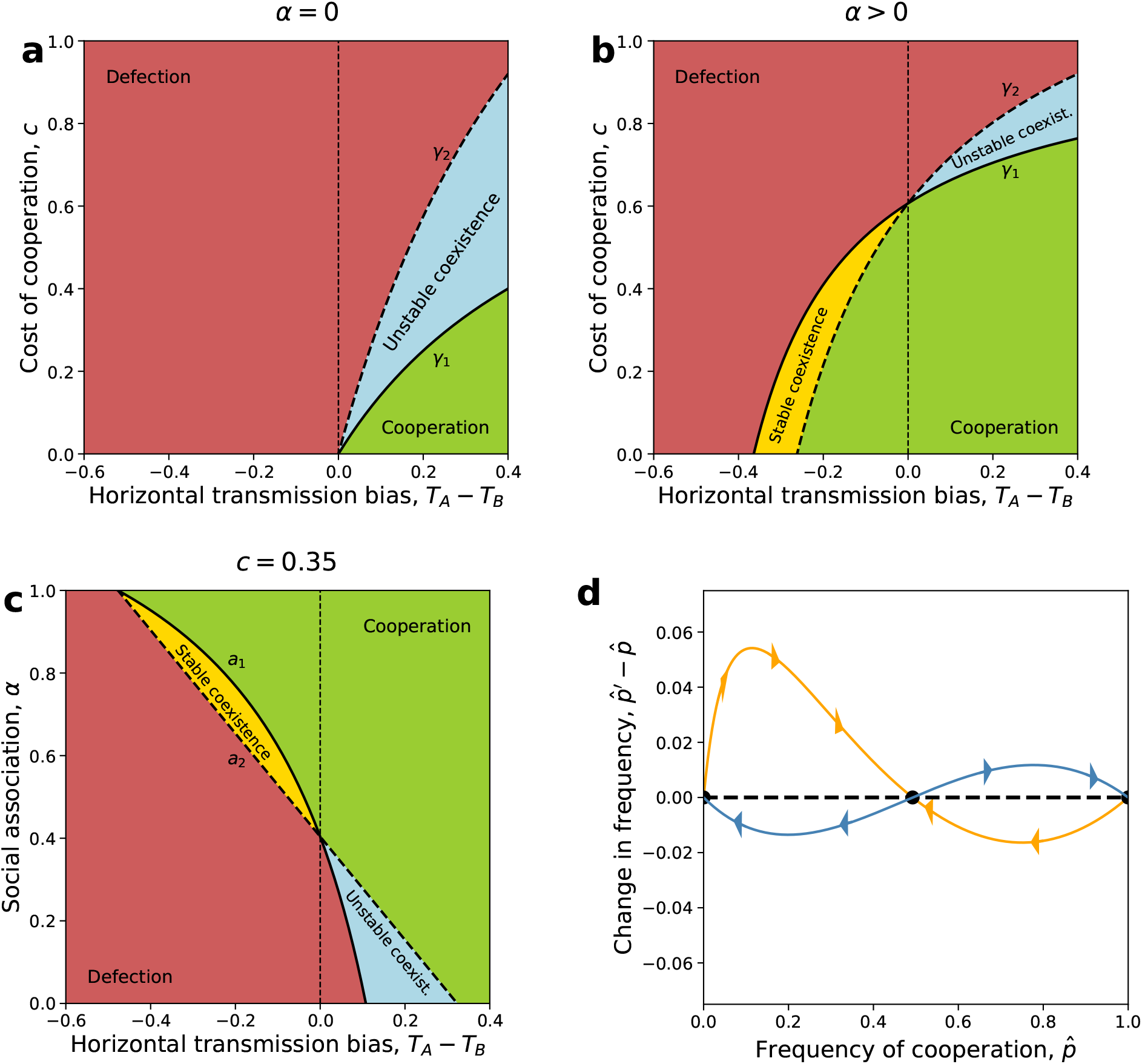
Evolution of cooperation under vertical and horizontal cultural transmission (*v*=1). The figure shows parameter ranges for global fixation of cooperation (green), global fixation of defection (red), fixation of either cooperation or defection depending on the initial conditions, i.e. unstable polymorphism (blue), and stable polymorphism of cooperation and defection (yellow). (**a-c**) The horizontal transmission bias (*T_A_* − *T_B_*) is on the x-axis. In panels **(a)** and **(b)**, the cost of cooperation *c* is on the y-axis and the cost thresholds *γ*_1_ and *γ*_2_ (Eq. 13) are the solid and dashed lines, respectively. In panel **(c)**, interaction-transmission association *α* is on the y-axis and the interaction-transmission association thresholds *a*_1_ and *a*_2_ (Eqs. 18) are the solid and dashed lines, respectively. Here, *b* = 1.3, *T_A_* = 0.4, *v* = 1, (a) *α* = 0, (b) *α* = 0.7, (c) *c* = 0.35. **(d)** Change in frequency of cooperation among juveniles 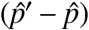 as a function of the frequency (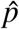), see Eq. 11. The orange curve shows convergence to a stable polymorphism (*T_A_* = 0.4, *T_B_* = 0.9, *b* = 12, *c* = 0.35, *v* = 1, and *α* = 0.45). The blue curve shows fixation of either cooperation or defection, depending on the initial frequency (*T_A_* = 0.5, *T_B_* = 0.1, *b* = 1.3, *c* = 0.904, *v* = 1, and *α* = 0.4). Black circles show the three equilibria.

Much of the literature on evolution of cooperation focuses on conditions for an initially rare cooperative phenotype to invade a population of defectors. The following remarks address this condition.

#### Remark 1 Condition on *c* for cooperation to increase when rare, i.e., 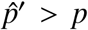 near *p* = 0

*If the initial frequency of cooperation is very close to zero, then its frequency will increase if the cost of cooperation is low enough,*

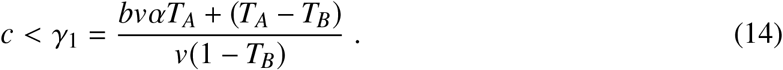

This unites the conditions for fixation of cooperation and for stable polymorphism, both of which entail instability of the state where defection is fixed, 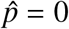.

Importantly, increasing interaction-transmission association *α* increases the cost threshold (∂*γ*_1_/∂*α* > 0), making it easier for cooperation to increase in frequency when initially rare. Similarly, increasing the horizontal transmission of cooperation, *T_A_*, increases the threshold (∂*γ*_1_/∂*T_A_* > 0), facilitating the evolution of cooperation ((Figure 3a and 3b). However, increasing the horizontal transmission of defection, *T_B_*, can increase or decrease the cost threshold, but it increases the cost threshold when the threshold is already above one (*c* < 1 < *γ*_1_): ∂*γ*_1_/∂*T_B_* is positive when 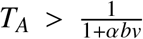, which gives *γ*_1_ > 1/*v*. Therefore, increasing *T_B_* decreases the cost threshold and limits the evolution of cooperation, but only if 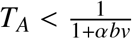.

Increasing the vertical transmission rate, *v*, can either increase or decrease the cost threshold, depending on the horizontal transmission bias, *T_A_* − *T_B_*, because sign (∂*γ*_1_/∂*v*) = −sign(*T_A_* − *T_B_*). When *T_A_* < *T_B_* we have ∂*γ*_1_/∂*v* > 0, and as the vertical transmission rate increases, the cost threshold increases, making it easier for cooperation to increase when rare (Figure 2b). In contrast, when *T_A_* > *T_B_* we get ∂*γ*_1_/∂*v* < 0, and therefore as the vertical transmission rate increases, the cost threshold decreases, making it harder for cooperation to increase when rare (Figure 2a).

In general, this condition cannot be formulated in the form of Hamilton’s rule due to the bias in horizontal transmission, represented by *T_A_* − *T_B_*. When there is no horizontal transmission bias, *T_A_* = *T_B_*, the following applies.

From Result 1 and inequality 14, if horizontal transmission is unbiased, *T* = *T_A_* = *T_B_*, then cooperation will take over the population from any initial frequency if the cost is low enough,

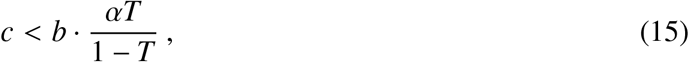

and regardless of the vertical transmission rate, *v*. This condition can be interpreted as a version of Hamilton’s rule (*c* < *b* · *r*, inequality 1) or as a version of inequality 3, where *αT*/(1 − *T*) can be regarded as the *effective relatedness* or *effective assortment*, respectively. Note that the right-hand side of inequality 15 equals *γ*_1_ when *T* = *T_A_* = *T_B_*.

From inequality 14, without interaction-transmission association (*α* = 0), cooperation will increase when it is rare if there is horizontal transmission bias for cooperation, *T_A_* > *T_B_*, and

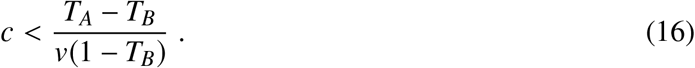

Figure 3a illustrates this condition (for *v* = 1), which is obtained by setting *α* = 0 in inequality 14. Importantly, the benefit of cooperation, *b*, does not affect the evolution of cooperation in the absence of interaction-transmission association, and the outcome is determined only by cultural transmission. Further, inequality 14 shows that with perfect interaction-transmission association (*α* = 1), cooperation will increase when rare if

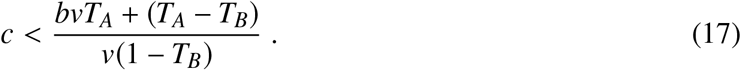

In the absence of oblique transmission, *v* = 1, the only equilibria are the fixation states, 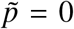 and 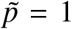, and cooperation will evolve from any initial frequency (i.e., 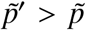) if inequality 17 applies (Figure 3). This is similar to case of microbe-induced cooperation studied by Lewin-Epstein *et al.* [16]; therefore when *v* = 1, this remark is equivalent to their eq. 1.

It is interesting to examine the general effect of interaction-transmission association *α* on the evolution of cooperation. Define the interaction-transmission association thresholds, *a*_1_ and *a*_2_, as

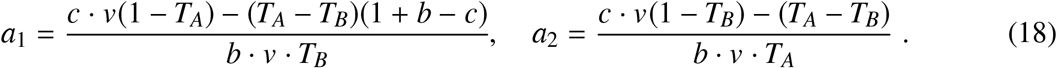

#### Remark 2 Condition on interaction-transmission association *α* for cooperation to increase when it is rare

*Cooperation will increase when rare if interaction-transmission association is high enough, specifically if a*_2_ < *α*.

Figures 2c and 2d illustrate this condition. With horizontal transmission bias for cooperation, *T_A_* > *T_B_*, cooperation can fix from any initial frequency if *a*_2_ < *α* (green area in the figures). With horizontal bias favoring defection, *T_A_* < *T_B_*, cooperation can fix from any frequency if interaction-transmission association is high, *a*_1_ < *α* (green area with *T_A_* < *T_B_*), and can also increase when rare and reach stable polymorphism if interaction-transmission association is intermediate, *a*_2_ < *α* < *a*_1_ (yellow area). Without horizontal bias, *T_A_* = *T_B_*, fixation of cooperation occurs if interaction-transmission association is high enough, 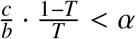 (inequality 15; in this case *a*_1_ = *a*_2_).

Interestingly, because sign (∂*a*_2_/∂*v*) = sign(*T_A_* − *T_B_*), the effect of the vertical transmission rate *v* on *a*_1_ and *a*_2_ depends on the horizontal transmission bias. That is, with horizontal bias for cooperation, *T_A_* > *T_B_*, evolution of cooperation is facilitated by oblique transmission, whereas with horizontal bias for defection, *T_A_* < *T_B_*, evolution of cooperation is facilitated by vertical transmission. This is demonstrated in Figures 2c and 2d.

Next, we examine the roles of vertical and oblique transmission in the evolution of cooperation. Fixation of cooperation is possible only if the vertical transmission rate is high enough,

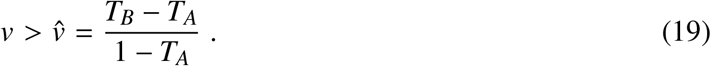

This condition is necessary for fixation of cooperation, but it is not sufficient. If horizontal transmission is biased for cooperation, *T_A_* > *T_B_*, cooperation can fix with any vertical transmission rate (because 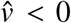). In contrast, if horizontal transmission is biased for defection, *T_A_* < *T_B_*, cooperation can fix only if the vertical transmission rate is high enough: in this case oblique transmission can prevent fixation of cooperation (see Figures 2b and 2d).

With only vertical transmission (*v* = 1), inequality 14 entails that cooperation will increase when rare if

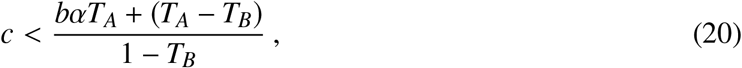

which can also be written as

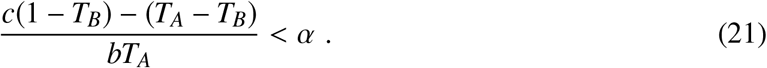

In the absence of vertical transmission (*v* = 0), from recursion 11 we see that the frequency of the cooperator phenotype among adults increases every generation, i.e. *p*′ > *p*, if there is a horizontal transmission bias in favor of cooperation, namely *T_A_* > *T_B_*. That is, in the absence of vertical transmission, selection plays no role in the evolution of cooperation (i.e., *b* and *c* do not affect 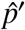). The dynamics are determined solely by differential horizontal transmission of the two phenotypes, namely, the relative tendency of each phenotype to be horizontally transmitted to peers. With no bias in horizontal transmission, *T_A_* = *T_B_*, phenotype frequencies do not change, 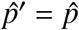.

Cooperation and defection can coexist at frequencies 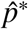 and 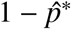 (Eq. 12). When it is feasible, this equilibrium is stable or unstable under the conditions of Result 1, parts 3 and 4, respectively. The yellow and blue areas in Figures 3 and 2 show cases of stable and unstable polymorphism, respectively. When 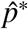 is unstable, cooperation will fix if its initial frequency is 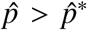, and defection will fix if 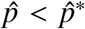. 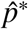 is unstable when there is horizontal transmission bias for cooperation, *T_A_* > *T_B_*, and the cost is intermediate, *γ*_1_ < *c* < *γ*_2_. Figure 3d shows 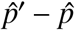 as a function of 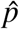.

### Evolution of interaction-transmission association

We now focus on the evolution of interaction-transmission association under perfect vertical transmission, *v* = 1, assuming that the population is initially at a stable polymorphism of the two phenotypes, cooperation *A* and defection *B*, where the frequency of *A* among juveniles is 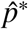 (Eq. 12). Note that for a stable polymorphism, there must be horizontal bias for defection, *T_A_* < *T_B_*, and an intermediate cost of cooperation, *γ*_2_ < *c* < *γ*_1_ (Eq. 13), see Figure 3b. The equilibrium population mean fitness is 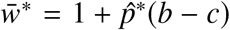, which is increasing in 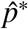, and 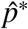 is increasing in *α* (Appendix C). Therefore, if interaction-transmission association increases, the population mean fitness also increases. But can this population-level advantage lead to the evolution of interaction-transmission association?

To answer this question, we extend our model to include a “modifier locus” [7, 18, 19, 20] that determines interaction-transmission association, but has no direct effect on fitness. The modifier locus has two alleles, *M* and *m*, which induce interaction-transmission associations *α*_1_ and *α*_2_, respectively. Suppose that the population has evolved to a stable equilibrium 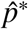 when only allele *M* is present. We study the local stability of this equilibrium to invasion by the modifier allele *m*; this is called “external stability” [1, 20] and obtain the following result.

#### Result 2 Reduction principle for interaction-transmission association

*From a stable polymorphism between cooperation and defection, a modifier allele can successfully invade the population if it decreases the interaction-transmission association α*.

The full analysis is in Appendix D. Note that this reduction principle entails that successful invasions will reduce the frequency of cooperation, as well as the population mean fitness (Figure 4). Furthermore, if we assume that modifier alleles with decreased interaction-transmission association appear and invade the population from time to time, then interaction-transmission association will continue to decrease, further reducing the frequency of cooperation and the population mean fitness. This evolution will proceed as long as there is a stable polymorphism, that is, as long as *a*_2_ < *α* < *a*_1_ (Remark 2, Figure 3c). Thus, we can expect interaction-transmission association to eventually approach *a*_2_, the frequency of cooperation to fall to zero, and the population mean fitness to decrease to one (Figure 4).

**Figure 4:**
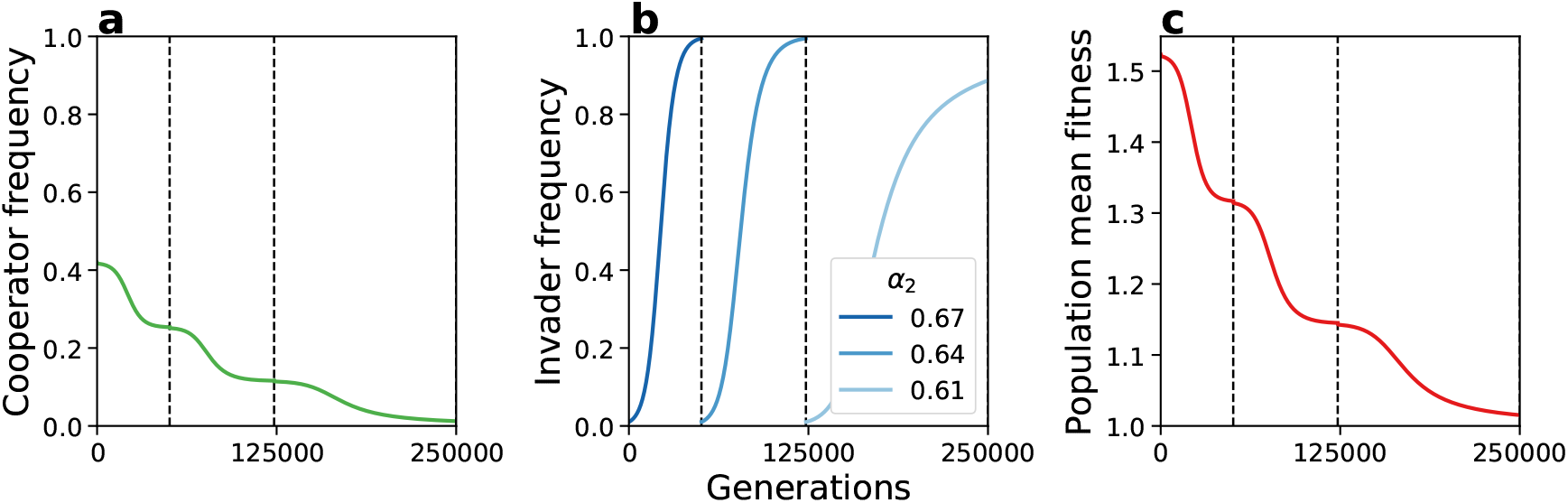
Reduction principle for interaction-transmission association. Consecutive fixation of modifier alleles that reduce interaction-transmission association *α* in numerical simulations of evolution with two modifier alleles (Eq. D1). When an invading modifier allele is established in the population (frequency > 99.95%), a new modifier allele that reduces interaction-transmission association by 5% is introduced (at initial frequency 0.5%). **(a)** The frequency of the cooperative phenotype *A* over time. **(b)** The frequency of the invading modifier allele *m* over time. **(c)** The population mean fitness 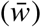 over time. Here, *c* = 0.05, *b* = 1.3, *T_A_* = 0.4 < *T_B_* = 0.7, initial interaction-transmission association *α*_1_ = 0.7, lower interaction-transmission association threshold *a*_2_ = 0.605.

### Population structure

Interaction-transmission association may also emerge from population structure. Consider a population colonizing a two-dimensional grid of size 100-by-100, where each site is inhabited by one individual, similarly to the model of Lewin-Epstein & Hadany [17]. Each individual is characterized by its phenotype: either cooperator, *A*, or defector, *B*. Initially, each site in the grid is randomly colonized by either a cooperator or a defector, with equal probability. In each generation, half of the individuals are randomly chosen to “initiate” interactions, and these initiators interact with a random neighbor (i.e. individual in a neighboring site) in a prisoners’ dilemma game (Figure 1a) and a random neighbor (with replacement) for horizontal cultural transmission (Figure 1b). The expected number of each of these interactions per individual per generation is one. The effective interaction-transmission association *α* in this model is the probability that the same neighbor is picked for both interactions, or *α* = 1/*M*, where *M* is the number of neighbors. On an infinite grid, *M* = 8, but on a finite grid *M* can be lower in edge neighborhoods close to the grid border. As before, *T_A_* and *T_B_* are the probabilities of successful horizontal transmission of phenotypes *A* and *B*, respectively.

The order of the interactions across the grid at each generation is random. After all interactions take place, an individual’s fitness is determined by *w* = 1+*b*·*n_b_*−*c*·*n_c_*, where *n_b_* is the number of interactions that individual had with cooperative neighbors, and *n_c_* is the number of interactions in which that individual cooperated (note that the phenotype may change between consecutive interactions due to horizontal transmission). Then, a new generation is produced, and the sites can be settled by offspring of any parent, not just the neighboring parents. Selection is global, rather than local, in accordance with our deterministic model: The parent is randomly drawn with probability proportional to its fitness, divided by the sum of the fitness values of all potential parents. Offspring are assumed to have the same phenotype as their parents (i.e. *v* = 1).

The outcomes of stochastic simulations with such a structured population are shown in Figure 5, which demonstrates that the highest cost of cooperation *c* that permits the evolution of cooperation agrees with the conditions derived above for our model without population structure or stochasticity. An example of stochastic stable polymorphism is shown in Figure 5c. Changing the simulation so that selection is local (i.e., sites can only be settled by offspring of neighboring parents) had only a minor effect on the agreement with the derived conditions (Figure S1).

**Figure 5:**
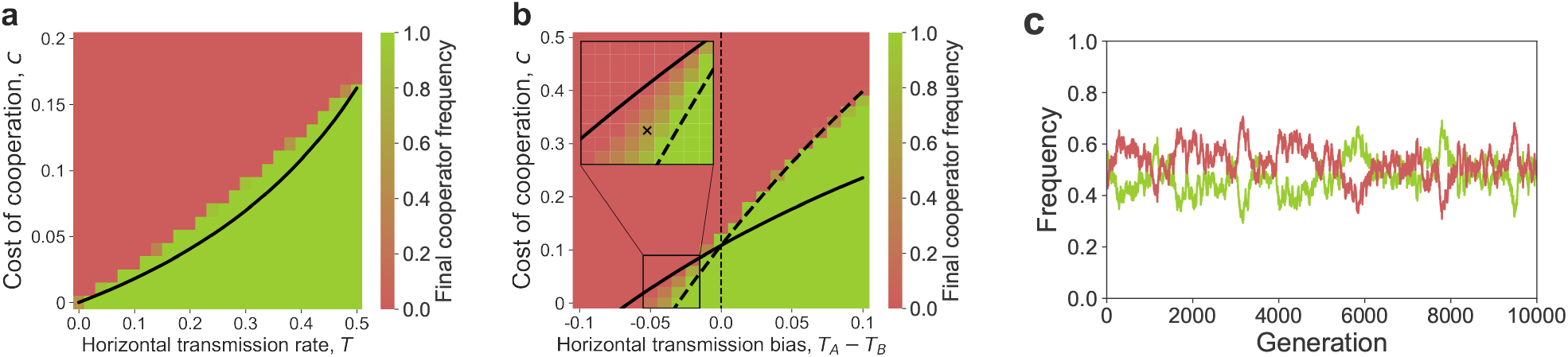
Evolution of cooperation in a structured population. **(a-b)** The expected frequency of cooperators in a structured population after 10,000 generations is shown (red for 0%, green for 100%) as a function of both the cost of cooperation, *c*, on the y-axis, and either the symmetric horizontal transmission rate, *T* = *T_A_* = *T_B_*, on the x-axis of panel **(a)**, or the transmission bias, *T_A_* − *T_B_*, on the x-axis of panel **(b)**. Black curves represent the cost thresholds for the evolution of cooperation in a well-mixed population with interaction-transmission association, where *α* = 1 8 in inequality 15 for panel **(a)** and in Eqs. 13 for panel **(b)**. The inset in panel **(b)** focuses on an area of the parameter range in which neither phenotype is fixed throughout the simulation, maintaining a stochastic locally stable polymorphism [14]. This stochastic polymorphism is illustrated in panel **(c)**, which shows the frequency of cooperators (green) and defectors (red) over time for the parameter set marked by an *x* in panel **(b)**. In all cases, the population evolves on a 100-by-100 grid. Cooperation and horizontal transmission are both local between neighboring sites, and each site has 8 neighbors. Selection operates globally (see Figure S1 for results from a model with local selection). Simulations were stopped at generation 10,000 or if one of the phenotypes fixed. 50 simulations were executed for each parameter set. Benefit of cooperation, *b* = 1.3; perfect vertical transmission *v* = 1. **(a)** Symmetric horizontal transmission, *T* = *T_A_* = *T_B_*; **(b)** Horizontal transmission rate *T_A_* is fixed at 0.4, and *T_B_* varies, 0.3 < *T_B_* < 0.5. **(c)** Horizontal transmission rates *T_A_* = 0.4 < *T_B_* = 0.435 and cost of cooperation *c* = 0.02.

These comparisons between the deterministic unstructured model and the stochastic structured model show that the conditions derived for the deterministic model can be useful for predicting the dynamics under complex scenarios. Moreover, this structured population model demonstrates that our parameter for interaction-transmission association, *α*, can represent local interactions between individuals.

## Discussion

Under a combination of vertical, oblique, and horizontal transmission with payoffs in the form of a prisoner’s dilemma game, cooperation or defection can either fix or coexist, depending on the relationship between the cost and benefit of cooperation, the horizontal transmission bias, and the association between social interaction and horizontal transmission (Result 1, Figures 2 and 3). Importantly, cooperation can increase when initially rare (i.e. invade a population of defectors) if and only if, rewriting inequality 14,

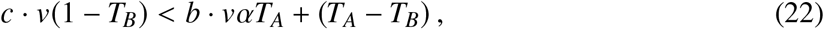

namely, the effective cost of cooperation (left-hand side) is smaller then the effective benefit plus the horizontal transmission bias (right-hand side). This condition cannot be formulated in the form of Hamilton’s rule, *c* < *b* · *r*, due to the effect of biased horizontal transmission, represented by *T_A_* − *T_B_*. Remarkably, a polymorphism of cooperation and defection can be stable if horizontal transmission is biased in favor of defection (*T_A_* < *T_B_*) and both the cost of cooperation and interaction-transmission association are intermediate (yellow areas in Figures 2 and 3).

We find that stronger interaction-transmission association *α* leads to evolution of higher frequency of cooperation and increased population mean fitness. Nevertheless, when cooperation and defection coexist, interaction-transmission association is expected to be reduced by natural selection, leading to extinction of cooperation and decreased population mean fitness (Result 2, Figure 4). Without interaction-transmission association, the benefit of cooperation cannot facilitate its evolution, and cooperation can only succeed if horizontal transmission is biased in its favor.

Indeed, in our model, horizontal transmission plays a major role in the evolution of cooperation: increasing the transmission of cooperation, *T_A_*, or decreasing the transmission of defection, *T_B_*, facilitates the evolution of cooperation. However, the effect of oblique transmission is more complicated. When there is horizontal transmission bias in favor of cooperation, *T_A_* > *T_B_*, increasing the rate of oblique transmission, 1 − *v*, will facilitate the evolution of cooperation. In contrast, when the bias is in favor of defection, *T_A_* < *T_B_*, higher rates of vertical transmission, *v*, are advantageous for cooperation, and the rate of vertical transmission must be high enough 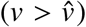 for cooperation to fix in the population.

The conditions derived from our deterministic model provide a good approximation to outcomes of simulations of a complex stochastic model with population structure in which individuals can only interact with and transmit to their neighbors. In these structured populations interaction-transmission association arises due to both social interactions and horizontal cultural transmission being local (Figure 5).

Feldman *et al.* [8] studied the dynamics of an altruistic phenotype with vertical cultural transmission and a gene that modifies the transmission of the phenotype. Their results are very sensitive to this genetic modification: without it, the conditions for invasion of the altruistic phenotype reduce to Hamilton’s rule. Further work is needed to incorporate such genetic modification of cultural transmission into our model. Woodcock [28] stressed the significance of non-vertical transmission for the evolution of cooperation and carried out simulations with prisoner’s dilemma payoffs but without horizontal transmission or interaction-transmission association (*α* = 0). Nevertheless, his results demonstrated that it is possible to sustain altruistic behavior via cultural transmission for a substantial length of time. He further hypothesized that horizontal transmission can play an important role in the evolution of cooperation, and our results provide strong evidence for this hypothesis.

To understand the role of horizontal transmission, we first review the role of *assortment*. Eshel & Cavalli-Sforza [6] showed that altruism can evolve when the tendency for *assortative meeting*, i.e., for individuals to interact with others of their own phenotype, is strong enough. Fletcher & Doebeli further argued that a general explanation for the evolution of altruism is given by *assortment*: the correlation between individuals that carry an altruistic trait and the amount of altruistic behavior in their interaction group (see also Bijma & Aanen [3]). They suggested that to explain the evolution of altruism, we should seek mechanisms that generate assortment, such as population structure, repeated interactions, and individual recognition. Our results highlight another mechanism for generating assortment: an association between social interactions and horizontal transmission that creates a correlation between one’s partner for interaction and the partner for transmission. This mechanism does not require repeated interactions, population structure, or individual recognition. We show that high levels of such interaction-transmission association greatly increase the potential for evolution of cooperation. With enough interaction-transmission association, cooperation can increase in frequency when initially rare even when there is horizontal transmission bias against it (*T_A_* < *T_B_*).

How does non-vertical transmission generate assortment? Lewin-Epstein *et al.* [16] and Lewin-Epstein & Hadany [17] suggested that microbes that induce their hosts to act altruistically can be favored by selection, which may help to explain the evolution of cooperation. From the kin selection point-of-view, if microbes can be transmitted *horizontally* from one host to another during host interactions, then following horizontal transmission the recipient host will carry microbes that are closely related to those of the donor host, even when the two hosts are (genetically) unrelated. From the assortment point-of-view, infection by behavior-determining microbes during interactions effectively generates assortment because a recipient of help may be infected by a behavior-determining microbe and consequently become a helper. Cultural horizontal transmission can similarly generate assortment between cooperators and enhance the benefit of cooperation if cultural transmission and helping interactions occur between the same individuals, i.e. when there is interaction-transmission association, so that the recipient of help may also be the recipient of the cultural trait for cooperation. Thus, with horizontal transmission, “assortment between focal cooperative players and cooperative acts in their interaction environment” [9] is generated not because the helper is likely to be helped, but rather because the helped is likely to become a helper.

## Acknowledgements

We thank Lilach Hadany, Ayelet Shavit, and Kaleda Krebs Denton for discussions and comments. This work was supported in part by the Clore Foundation Scholars Programme (OLE), the Morrison Institute for Population and Resources Studies at Stanford University (MWF), Israel Science Foundation (YR 552/19), and Minerva Stiftung Center for Lab Evolution (YR).

## Appendices

## Appendix A Local stability criterion

Let *f* (*p*) = *λ* · (*p*′ − *p*), where *λ* > 0, and 0 and 1 are equilibria, that is, *f* (0) = 0 and *f* (1) = 0.

Set *p* > *p*\* = 0. Using a linear approximation for *f* (*p*) near 0, we have

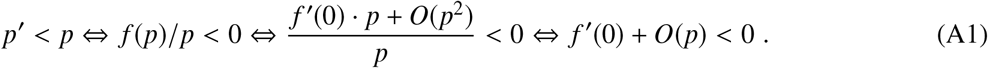

Therefore, by definition of big-O notation, if *f*′(0) < 0 then there exists *ϵ* > 0 such that for any local perturbation 0 < *p* < *ϵ*, it is guaranteed that 0 < *p*′ < *p*; that is, *p*′ is closer to zero than *p*.

Set *p* < *p*\* = 1 Using a linear approximation for *f*(*p*) near 1, we have

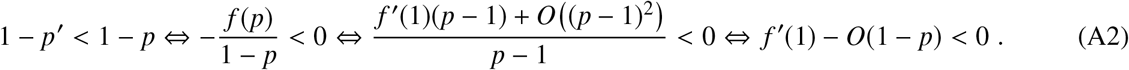

Therefore, if *f*′(1) < 0 then there exists *ϵ* > 0 such that for any 1 − *ϵ* < 1 − *p* < 1 we have 1 − *p*′ < 1 − *p*; that is, *p*′ is closer to one than *p*.

## Appendix B Equilibria and stability

Let 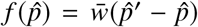. Then, using *SymPy* [22], a Python library for symbolic mathematics, this simplifies to

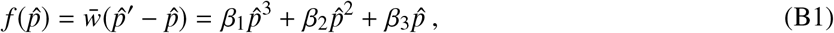

where

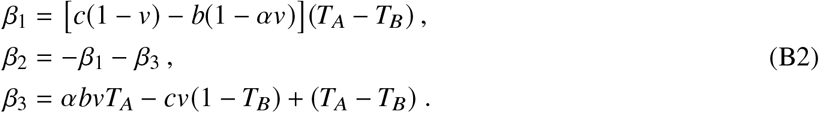

If *T* = *T_A_* = *T_B_* then *β*_1_ = 0 and *β*_3_ = −*β*_2_ = *αbvT* − *cv*(1 − *T*), and 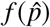 becomes a quadratic polynomial,

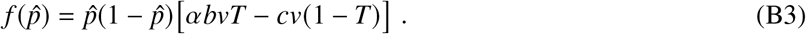

Clearly the only two equilibria are the fixations 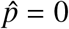 and 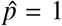, which are are locally stable if 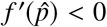 near the equilibrium (see Appendix A), where 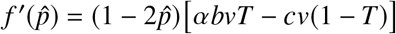, so that

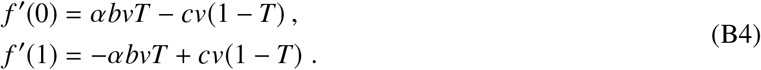

In the general case where *T_A_* ≠ *T_B_*, the coefficient *β*_1_ is not necessarily zero, and 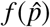 is a cubic polynomial. Therefore, three equilibria may exist, two of which are 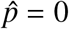 and 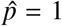, and the third is

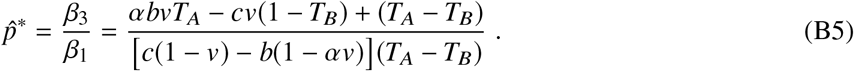

Note that the sign of the cubic (Eq. B1) at positive (negative) infinity is equal (opposite) to the sign of *β*_1_. If *T_A_* > *T_B_*, then

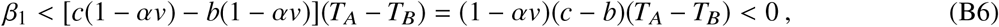

since *c* < *b* and *αv* < 1. Hence the signs of the cubic at positive and negative infinity are negative and positive, respectively. First, if *β*_3_ < *β*_1_ then 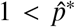. Also, *f*′(0) < 0 and *f*′(1) > 0; that is, fixation of the defector phenotype *B* is the only locally stable feasible equilibrium. Second, if *β*_1_ < *β*_3_ < 0 then 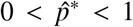 and therefore *f*′(0) < 0 and *f*′(1) < 0 so that both fixations are locally stable and 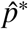 separates the domains of attraction. Third, if 0 < *β*_3_ then 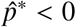 and therefore *f*′(0) > 0 and *f*′(1) < 0; that is, fixation of the cooperator phenotype *A* is the only locally stable legitimate equilibrium.

Similarly, if *T_A_* < *T_B_*, then

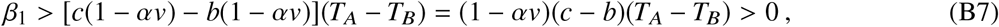

since *c* < *b* and *αv* < 1, and the signs of the cubic at positive and negative infinity are positive and negative, respectively. First, if *β*_3_ < 0 then 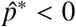 and therefore *f*′(0) < 0 and *f*′(1) > 0; that is, fixation of the defector phenotype *A* is the only locally stable legitimate equilibrium. Second, if 0 < *β*_3_ < *β*_1_ then 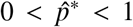 and therefore *f*′(0) > 0 and *f*′(1) > 0; that is, both fixations are locally unstable and 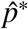 is a stable polymorphic equilibrium. Third, if *β*_1_ < *β*_3_ then 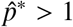 and therefore *f*′(0) > 0 and *f*′(1) < 0, and fixation of the cooperator phenotype *A* is the only locally stable feasible equilibrium.

This analysis can be summarized as follows:

1. *Fixation of cooperation*: if *(i) T* = *T_A_* = *T_B_* and 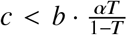; or if *(ii) T_A_* > *T_B_* and 0 < *β*_3_; or if *(iii) T_A_* < *T_B_* and *β*_1_ < *β*_3_.
2. *Fixation of the defection*: if *(iv) T* = *T_A_* = *T_B_* and 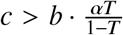; or if *(v) T_A_* > *T_B_* and *β*_3_ < *β*_1_ < 0; or if *(vi) T_A_* < *T_B_* and *β*_3_ < 0.
3. *polymorphism of both phenotypes at* 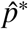: if *(vii) T_A_* < *T_B_* and 0 < *β*_3_ < *β*_1_.
4. *Fixation of either phenotype depending on initial frequency*: if *(viii) T_A_* > *T_B_* and *β*_1_ < *β*_3_ < 0.

We now proceed to use the cost thresholds, *γ*_1_ and *γ*_2_, and the vertical transmission threshold, 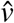 (Eq. 13). First, assume *T_A_* < *T_B_*. *β*_3_ < 0 requires *γ*_1_ < *c*. For *β*_3_ < *β*_1_ we need *c* [*v*(1 − *T_B_*) + (1 − *v*)(*T_A_* − *T_B_*)] > *bvαT_B_* + (1 + *b*)(*T_A_* − *T_B_*). Note that the expression in the square brackets is positive if and only if 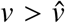. Thus, for *β*_3_ < *β*_1_ we need 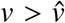 and *γ*_2_ < *c* or 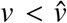 and *c* < *γ*_2_, and for 0 < *β*_3_ < *β*_1_ we need 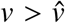 and *γ*_2_ < *c* < *γ*_1_, or 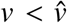 and *c* < min(*γ*_1_, *γ*_2_). For *β*_1_ < *β*_3_ we need 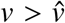 and *c* < *γ*_2_ or 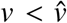 and *γ*_2_ < *c*. However, some of these conditions cannot be met, since 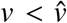 implies *c* < 1 < *γ*_2_.

Second, assume *T_A_* > *T_B_*. *β*_3_ > 0 requires *γ*_1_ > *c*. For *β*_1_ < *β*_3_ we need *c*[*v*(1 − *T_B_*) + (1 − *v*)(*T_A_* − *T_B_*)] < *bvαT_B_* + (1 + *b*)(T_A_ − *T_B_*). Thus for *β*_1_ < *β*_3_ we need 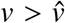 and *c* < *γ*_2_ or 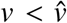 and *c* > *γ*_2_. But 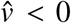 when *T_A_* > *T_B_*, and therefore we have *β*_1_ < *β*_3_ if *c* < *γ*_2_. Similarly, we have *β*_3_ < *β*_1_ if 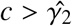.

This analysis is summarized in Result 1.

## Appendix C Effect of interaction-transmission association on mean fitness

To determine the effect of increasing *α* on the stable population mean fitness, 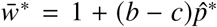, we must analyze its effect on 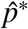,

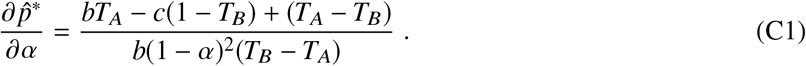

Note that stable polymorphism implies *c* < *γ*_1_, and because *α* < 1, we have

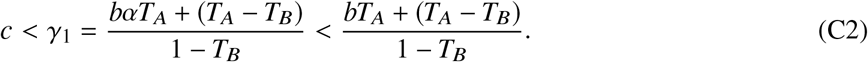

Therefore, the numerator in Eq. C1 is positive. Since *T_A_* < *T_B_*, the denominator in Eq. C1 is also positive, and hence the derivative 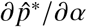 is positive. Thus, the population mean fitness increases as interaction-transmission association *α* increases.

## Appendix D Reduction principle

We assume here that *v* = 1, i.e. no oblique transmission, and therefore 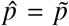. Denote the frequencies of the pheno-genotypes *AM*, *BM*, *Am*, and *Bm* by 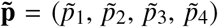. The frequencies of the pheno-genotypes in the next generation are defined by the recursion system,

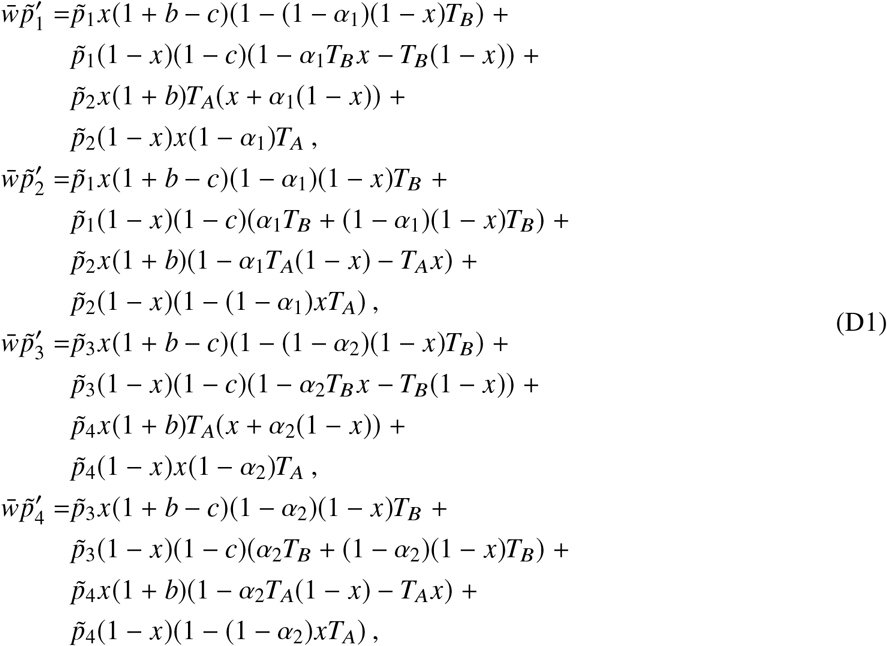

where 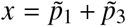 is the total frequency of the cooperative phenotype *A*, and 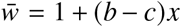 is the population mean fitness.

The equilibrium where only allele *M* is present is 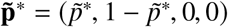, where

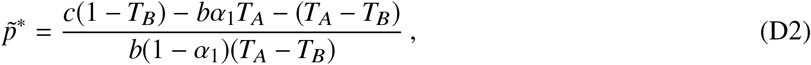

setting *α* = *α*_1_ and *v* = 1 in Eq. 12. When *v* = 1, 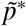 is a feasible polymorphism 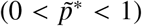 if *T_A_* < *T_B_* and *γ*_2_ < *c* < *γ*_1_ (Result 1).

The local stability of 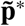 to the introduction of allele *m* is determined by the linear approximation **L**\* of the transformation in Eq. D1 near 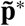 (i.e., the Jacobian of the transformation at the equilibrium). **L**\* is known to have a block structure, with the diagonal blocks occupied by the matrices 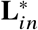 and 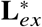 [1, 20]. The latter is the external stability matrix: the linear approximation to the transformation near 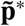 involving only the pheno-genotypes *Am* and *Bm*, derived from Eq. D1, with 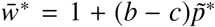 as the stable population mean fitness,

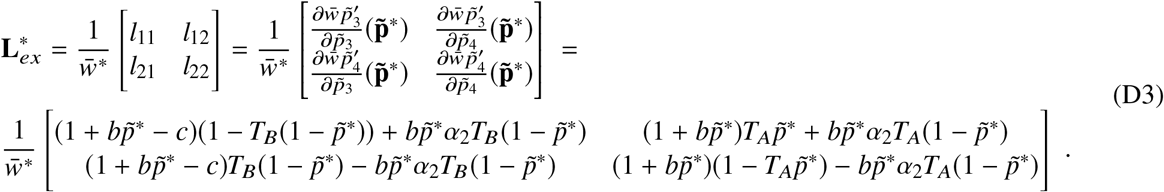

Because we assume that 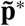 is internally stable (i.e. locally stable to small perturbations in the frequencies of *AM* and *BM*), the stability of 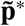 is determined by the eigenvalues of the external stability matrix 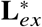. This is a positive matrix, and due to the Perron-Frobenius theorem, the leading eigenvalue of 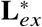 is real and positive. Thus, if the leading eigenvalue is less (greater) than one, then the equilibrium 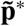 is externally stable (unstable) and allele *m* cannot (can) invade the population of allele *M*. The eigenvalues of 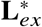 are the roots of the characteristic polynomial, *R*(*λ*), which is a quadratic with a positive leading coefficient. Therefore, lim_*λ*→±∞_ *R*(*λ*) = ∞, and the leading eigenvalue is less than one (implying stability) if and only if *R*(1) > 0 and *R*′(1) > 0. Thus, a sufficient condition for external instability of 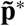 is *R*(1) < 0.

*R*(*λ*) is defined as a determinant, 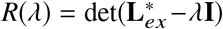, where **I** is the 2-by-2 identity matrix. Since multiplication by a positive factor doesn’t change the sign, and using the properties of the determinant, we have

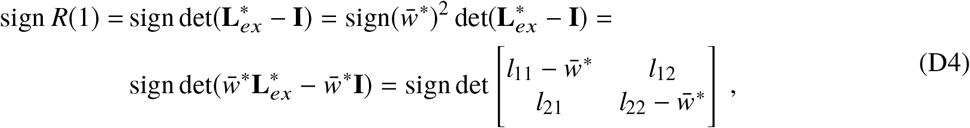

where *l_ij_* are defined in Eq. D3. Adding the second row in Eq. D4 to the first row, which does not change the determinant, and substituting 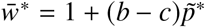, we get

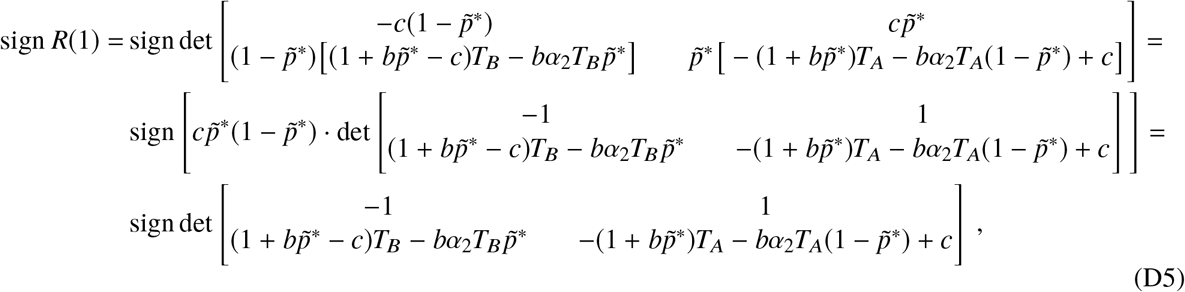

since *c* > 0, 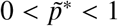. That is,

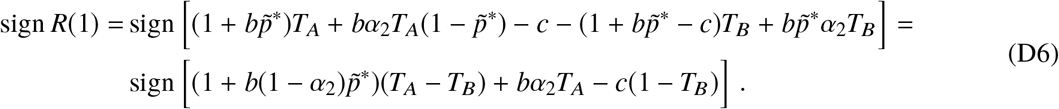

Substituting 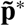 from Eq. D2, we get

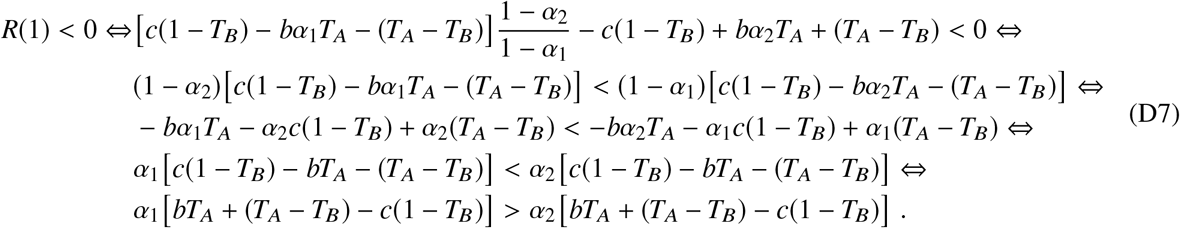

We assumed *c* < *γ*_1_, and since 0 ≤ *α*_1_ ≤ 1,

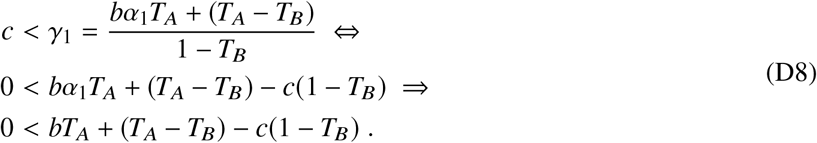

Combining inequalities D7 and D8, we find that *R*(1) < 0 if and only if *α*_1_ > *α*_2_, which is a sufficient condition for external instability. Therefore, if *α*_2_, the interaction-transmission association of the invading modifier allele *m*, is less than *α*_1_, the interaction-transmission association of the resident allele *M*, then invasion will be successful.

Determining a necessary and sufficient condition for successful invasion is more complicated, requiring analysis of the sign of *R*′(1). However, we have numerically validated that the leading eigenvalue is greater than one if and only if *α*_1_ > *α*_2_.

## Figures

**Figure S1:**
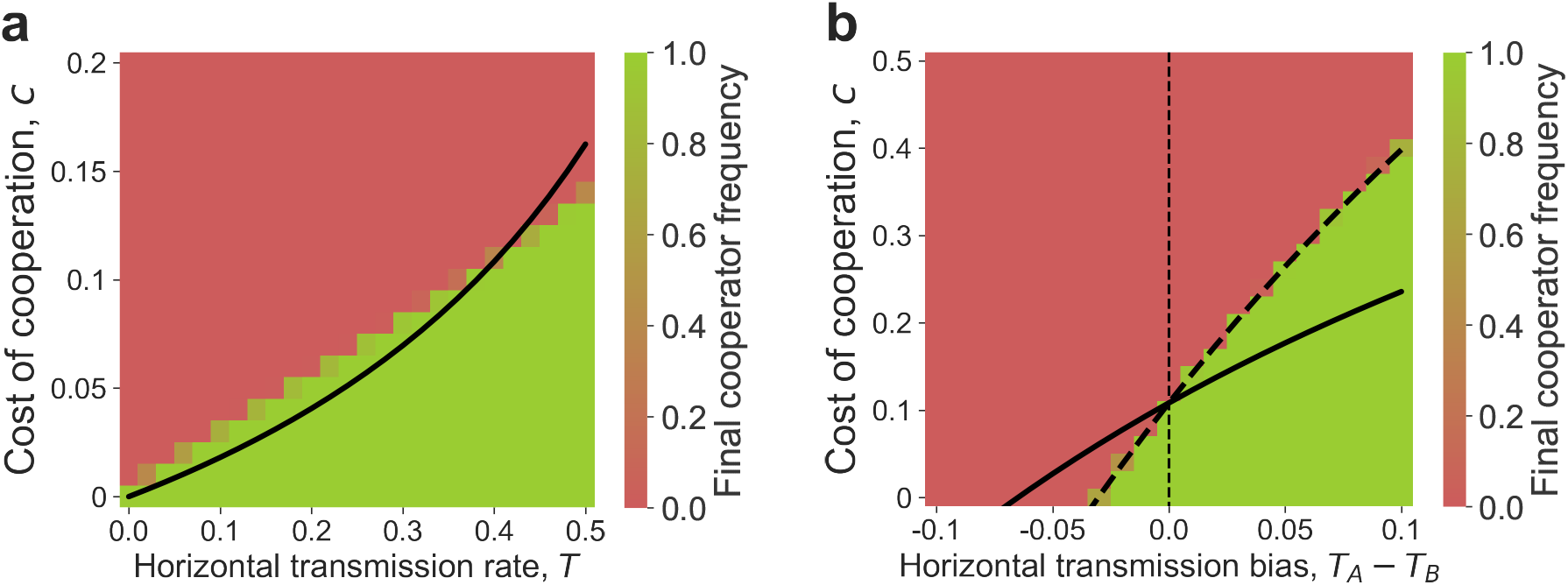
Evolution of cooperation in a structured population with local selection. The expected frequency of cooperators in a structured population after 10,000 generations is shown (red for 0%, green for 100%) as a function of both the cost of cooperation (*c*) on the y-axis, and the symmetric horizontal transmission rate (*T* = *T_A_* = *T_B_*) on the x-axis of panel **(a)**, or the transmission bias *T_A_* − *T_B_* on the x-axis of panel **(b)**. Cooperation and horizontal transmission are both local between neighboring sites, and each site had 8 neighbors. Selection operates locally (see Figure 5 for results from a model with global selection). The black curves represent the cost thresholds for the evolution of cooperation in a well-mixed population with interaction-transmission association, where *α* = 1 8 in inequality 15 for panel **(a)** and in Eqs. 13 for panel **(b)**. The population evolves on a 100-by-100 grid. Simulations were stopped at generation 10,000 or if one of the phenotypes fixed. 50 simulations were executed for each parameter set. Here, benefit of cooperation, *b* = 1.3; perfect vertical transmission *v* = 1. **(a)** Symmetric horizontal transmission, *T* = *T_A_* = *T_B_*. **(b)** Horizontal transmission rate *T_A_* is fixed at 0.4, and *T_B_* varies, 0.3 < *T_B_* < 0.5.

## Footnotes

1 In an extended model, which allows an individual to encounter *N* individuals before choosing a partner, the right hand side is multiplied by *E*[*N*], the expected number of encounters [6, eq. 4.6].

2 Inequality 3 generalizes inequalities 1 and 2 by substituting *p_C_* = *r* + *p*, *p_D_* = *p* and *p_C_* = *m* + (1 − *m*)*p*, *p_D_* = (1 − *m*)*p*, respectively, where *p* is the frequency of cooperators.

